# Large-scale analysis of transcript data reveals thousands of recursive splicing events in human introns

**DOI:** 10.64898/2026.09.10.750672

**Authors:** David J. Bass, Steven L. Salzberg

## Abstract

Recursive splicing (RS) is a process in which an intron is removed from a nascent RNA molecule in two or more splicing events rather than one. We introduce a novel approach for detecting recursive splice sites (RSSs), the intronic loci at which RS events occur, based on alignment of total RNA-seq data to short, customized “target” sequences. We applied this approach to a data set from a recent study of gene expression in the human brain, using parameters corresponding to a very low false discovery rate, and found 3,022 RSSs that appear in 2,775 distinct introns from 2,407 genes. 2,891 (96%) of these RSSs are in protein-coding genes. The median length of recursively spliced introns from this set is 10,114 base pairs, which is substantially longer than the median human intron, but much shorter than average RS intron lengths reported in prior studies. Our work dramatically increases the number of known RSSs in the human genome and provides a generalizable bioinformatics pipeline for annotating RSSs from total RNA-seq data.

**Author Summary:** RNA splicing is a necessary step in the processing of most eukaryotic RNA molecules. Sometimes splicing includes an intermediate step known as “recursive splicing,” through which introns are split into two subintrons, each of which is independently excised by the splicing machinery. Over the last decade, several studies have attempted to catalog recursive splice sites in humans and other model organisms, achieving little agreement across methods. While previous studies primarily used targeted laboratory methods, we sought to leverage previously generated large-scale human RNA-sequencing data sets to identify recursive splice sites. We specifically used a data set of total RNA-seq data, which contains many premature transcripts that are only partially spliced. We developed a method based on two parallel read alignment steps using both spliced and unspliced alignments of these reads. We found 3022 high-confidence recursive splice sites, which represents a 30-fold increase as compared to the largest prior curated set.

## Introduction

RNA splicing is a necessary and ubiquitous process in eukaryotic genes. Introns in precursor messenger RNA (pre-mRNA) must be excised via splicing. Recursive splicing (RS) is a non-canonical splicing mechanism in which a single intron is removed in multiple steps, as if it were actually two adjacent introns. In the years following the initial discovery of RS in *Drosophila melanogaster* (1), multiple groups have attempted to find and annotate human RS sites (2–6). Previously, Zhang *et al*. (4) characterized RS as relatively rare and non-constitutive, while Wan *et al*. (5) characterize recursive splice site (RSS) choice as a more common process that frequently selects non-canonical splice sites. Here we instead seek to identify and catalog frequently-used human RSSs using evidence from large-scale RNA-sequencing experiments.

Although hundreds of RS introns have been catalogued in *D. melanogaster* (7) and other vertebrates (3), relatively few have been catalogued in humans. Among those identified in humans, the scientific community remains far from agreement. For example, Hoppe *et al*. (6) report that their set of 94 RS introns had no overlap with smaller sets of RS introns reported in three preceding studies (2–4). This notable lack of consensus motivated our development of a new method for RSS detection.

Two of the clearest forms of evidence of recursive splicing available via high-throughput transcriptome sequencing are (i) sawtooth patterns in read coverage and (ii) direct lariat sequencing (8). The sawtooth pattern is a signature in the read coverage of an RNA-Seq experiment in which coverage decreases linearly in the 5′ to 3′ direction along an intron, and then rises again at the beginning of each exon. This pattern arises in total RNA-seq data (but not in poly-A RNA-seq data) as a side effect of co-transcriptional splicing, in which introns are removed soon after their 3′ end is transcribed (9). As a result, the 5′ portion of an intron is more likely to be sampled in an RNA-seq experiment, causing coverage to decrease linearly along the intron. Lariat sequencing (ii) relies on the phenomenon whereby an intron forms a “lariat” configuration when its 5′ end is briefly bound to a branch point located some 18–35 nucleotides from the 3′ end (10), creating a short-lived, free-floating lariat molecule. Reads that cross the lariat boundary can be identified by both the discontinuous (in the genome) jump from the 5′ end of the intron to the branchpoint and by a common transcription error that reverse transcriptase makes at the branchpoint. Pineda and Bradley (11) pioneered a method for lariat selection from standard RNA-Seq experiments, which was used by Hoppe *et al*. (6) to filter trillions of reads from the NCBI Sequence Read Archive, yielding a tiny proportion that originated from lariats.

While lariat reads provide the most direct evidence of a splicing event, they are expensive to extract. In contrast, our method detects RS directly from bulk total RNA-seq data, lowering both the computational and experimental costs of RS annotation.

With their lariat-based method, Hoppe *et al*. (6) reported finding 100 recursive splice sites (RSSs) within 94 human introns. Zhang *et al*. (4) used 4sUDRB-Seq to detect 342 unique “candidate” RSSs across three tissues, 19 of which exhibited a sawtooth pattern in RNA-Seq read coverage. They argued that RS is tissue-specific and non-constitutive; i.e., it is often replaced by standard splicing that removes the whole intron in a single step. Sibley *et al*. (3) used both RNA-Seq and iCLIP coverage to detect 434 putative RSSs, which they filtered down to just 9 finalist RS introns. Wan *et al*. 2021 performed RNA-seq between pulses of a pulse-chase experiment to report 5,983 RSSs, but only 27% (1,589) of their sites contained the `AG/GT` canonical acceptor–donor motif that other methods, including our own, require as a criterion for RSS detection.

While there is no consensus relationship between RS and human disease, numerous studies have linked splicing anomalies to disease; e.g., intron retention in brain cells has been linked to Alzheimer’s disease (12). Annotating human RSSs is an important piece in the puzzle of determining the causes and implications of pathogenic splicing defects. We note that the RSSs reported here should not be conflated with cryptic splice sites, which have been associated (for example) with ALS–FTD (13); none of our RSSs lie within 50 bp of the 41 cryptic exons annotated by Ling et al. Cataloging all modes of RNA processing, including recursive splicing, enhances our understanding of the splicing process and may allow us to re-interpret RNA-seq reads that were previously discarded as transcriptional noise or technical artifacts. Along with our predicted RSSs, we release our analysis pipeline as open-source software to be used to investigate RS in other RNA-seq data sets from humans and other species.

## Results

### A novel annotation of 3,022 human RSSs

We developed a computational method that aligns total RNA-seq data to both the subsequences of the nascent transcriptome and the genome in order to detect RSSs. We used data from 900 samples (see Methods) published in an earlier study from the Lieber Institute for Brain Development (LIBD) (14), applying our method using two annotation databases, CHESS (15) and GENCODE (16), to define the transcriptome. We found 3,187 RSS “semifinalists” from our alignments to target subsequences from the nascent transcriptome, which we narrowed down to 3,022 RSS finalists by corroborating them using spliced alignments to the genome.

Starting with bulk total RNA-seq reads and human genome annotations, we developed a pipeline (**Fig. 1a**) to collect evidence for RSSs from alignments to genomic DNA, spliced RNA, and customized target sequences. We combined information from the CHESS and GENCODE databases, from scores assigned to annotated introns by SPLAM (17), and from alignments of total RNA-Seq reads in order to obtain a set of pre-RNA targets, each of which represented a potential RS event. All candidate RS sites had to contain the sequence `AGGT`, a motif that contains the canonical acceptor dinucleotide `AG` immediately followed by canonical donor dinucleotide `GT`.

**Figure 1:**
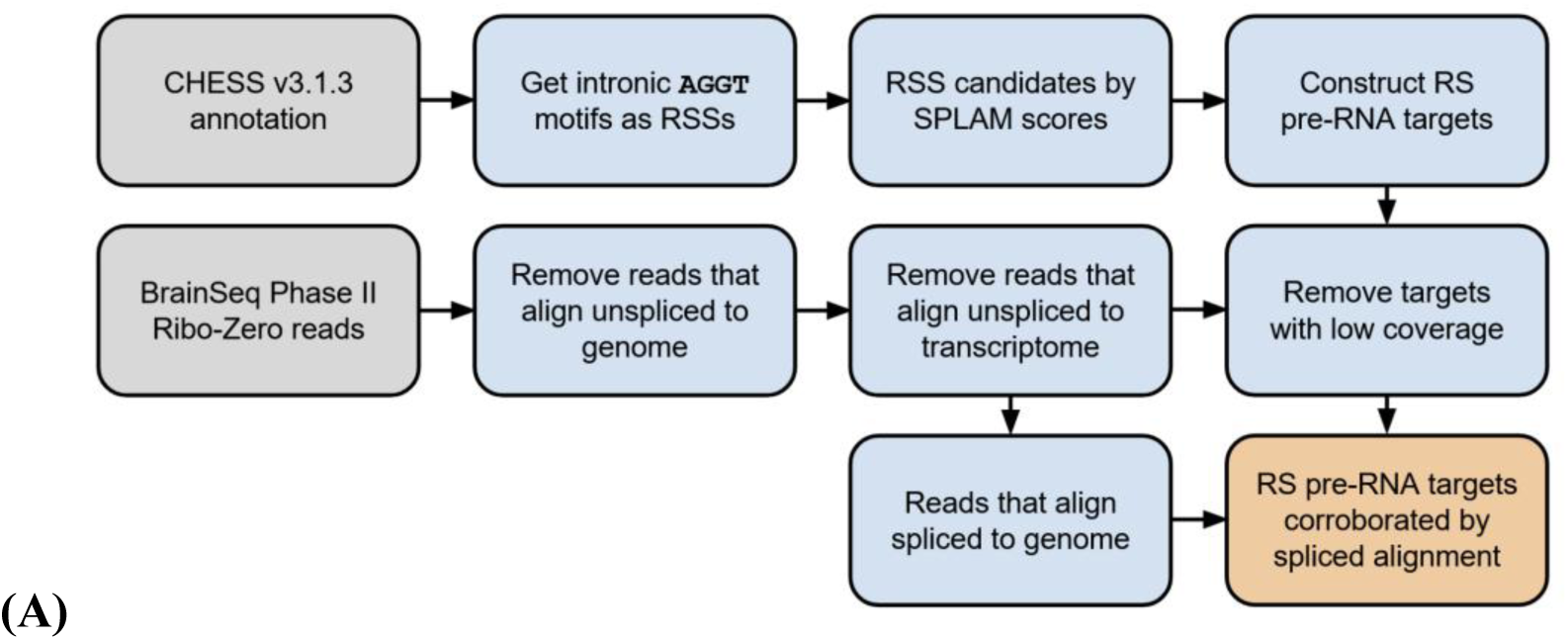
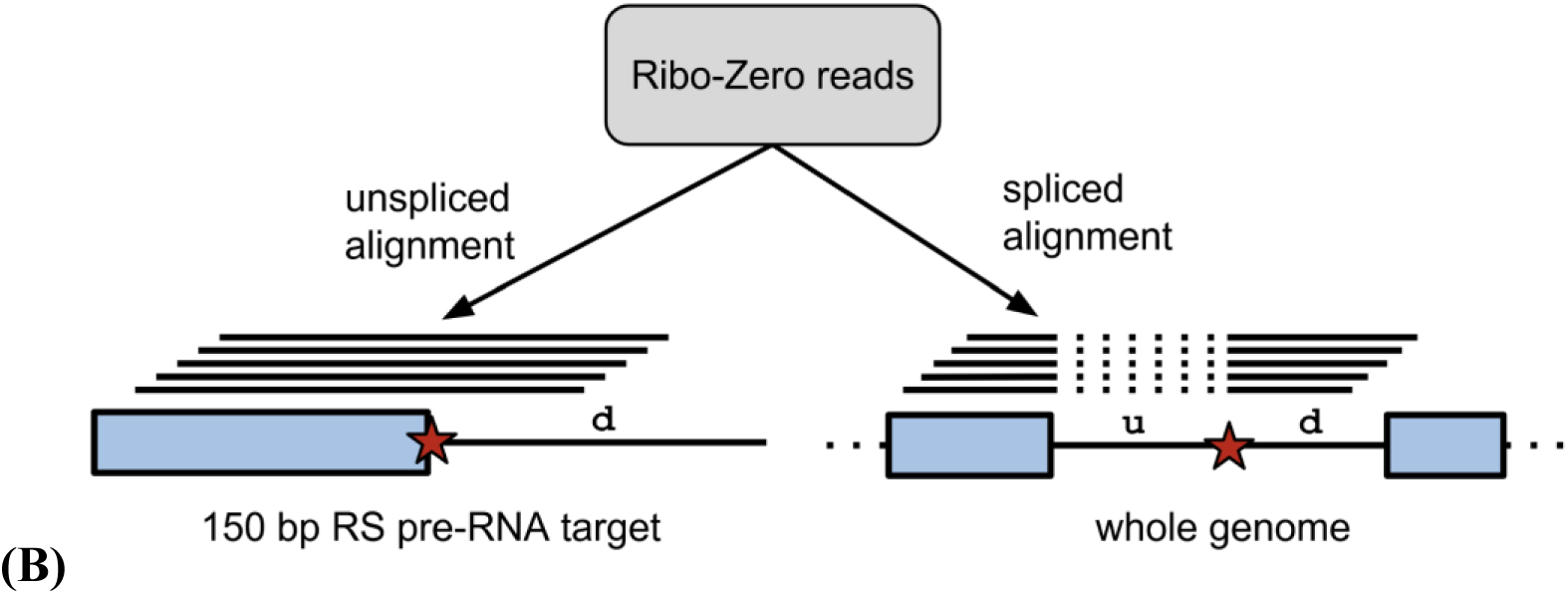
**(A)** Outline of our recursive splice site detection pipeline. **(B)** Illustration of a Ribo-Zero read pileup to alignment targets (left) using unspliced (Bowtie2) alignments to the genome, and (right) using spliced (HISAT2) alignments. `u` and `d` denote upstream and downstream subintrons respectively. The red star marks the recursive splice site, constrained to be centered on the sequence `AGGT`. Dotted lines in the pileup on the right denote the intron identified by the spliced alignment.

To build our customized alignment targets, we concatenated the last 75 bp of an exon with a 75 bp sequence that started at the `GT` of an `AGGT` tetranucleotide in each candidate RS site (**Fig. 1b**). This generated a set of 150bp target sequences that could only arise from recursive splicing that removed an upstream subintron and retained the downstream subintron; thus we could infer that reads aligning to each target were derived from RS events. Separately, we aligned the same read set to the entire genome, under the expectation that true RS events would be corroborated by reads that aligned in a spliced manner to the same location. Ideally, each read that aligned to an RS target would also align (spliced) over the corresponding subintron in the genome.

We began our search for RSS candidates with genomic evidence. We first identified all instances of the canonical RS motifs `AGGT` that occurred ≥50 bp from either end of an annotated intron of length ≥200. CHESS v3.1.3 contains 10,574,616 such loci. These thresholds eliminated candidate RS introns that were likely too short for RS to be sterically favorable. Second, we filtered the results to retain only those loci that received scores ≥0.5 from SPLAM, a deep-learning program for predicting splice sites (17). 1,674,568 loci (16%) passed this filter. While our discussion below focuses on the CHESS annotation, we replicated all analyses using the GENCODE annotation and we report those results as well.

We further filtered this candidate RSS set using transcriptomic evidence. Using a negative control set of 100,000 random loci, we set stringent thresholds to filter our candidates as follows: (i) ≥40 reads aligned to the target at (ii) ≥23 (of 51 possible) distinct alignment positions in the 150 bp target, and (iii) the reads came from ≥28 (of 900) Ribo-Zero samples. These three coverage metrics ensured that (i) multiple reads in the data set supported the RS event, (ii) these reads did not align in an aberrantly “clumpy” pattern in the target sequence, and (iii) these reads originated from many different biological samples. Only 9 targets from our set of 100,000 negative controls passed all three of these filters, yielding an expected false discovery rate of approximately one in 10,000.

From our 900 brain samples, these filtering steps yielded 3,187 RSSs that we labeled as semifinalists. We then removed 165 RSSs that had no support from the spliced alignments of the RNA-seq data set to the whole genome, yielding our final set of 3,022 RSSs. Based on the negative controls, the expected number of false positives in this set is less than one.

**Fig. 2B** shows that these findings did not show any obvious tissue-specific bias; we found 2,431 (80%) of the finalist RSSs when we used samples from just one tissue (either dorsolateral pre-frontal cortex (DLPFC) or hippocampus) or both tissues. Limiting our analysis to only one tissue slightly reduced the number of RSSs (2,768 from DLPFC, 2,923 from hippocampus). **Fig. 2C** shows that the RSSs are roughly uniformly distributed within their introns, with a slight bias towards the edges of introns. **Fig. 2D** shows that our finalists display the polypyrimidine tract found in most acceptor splice sites, with 86.7% of our RSS tetranucleotide motifs containing an upstream pyrimidine that matches the `YAGGT` motif.

**Figure 2:**
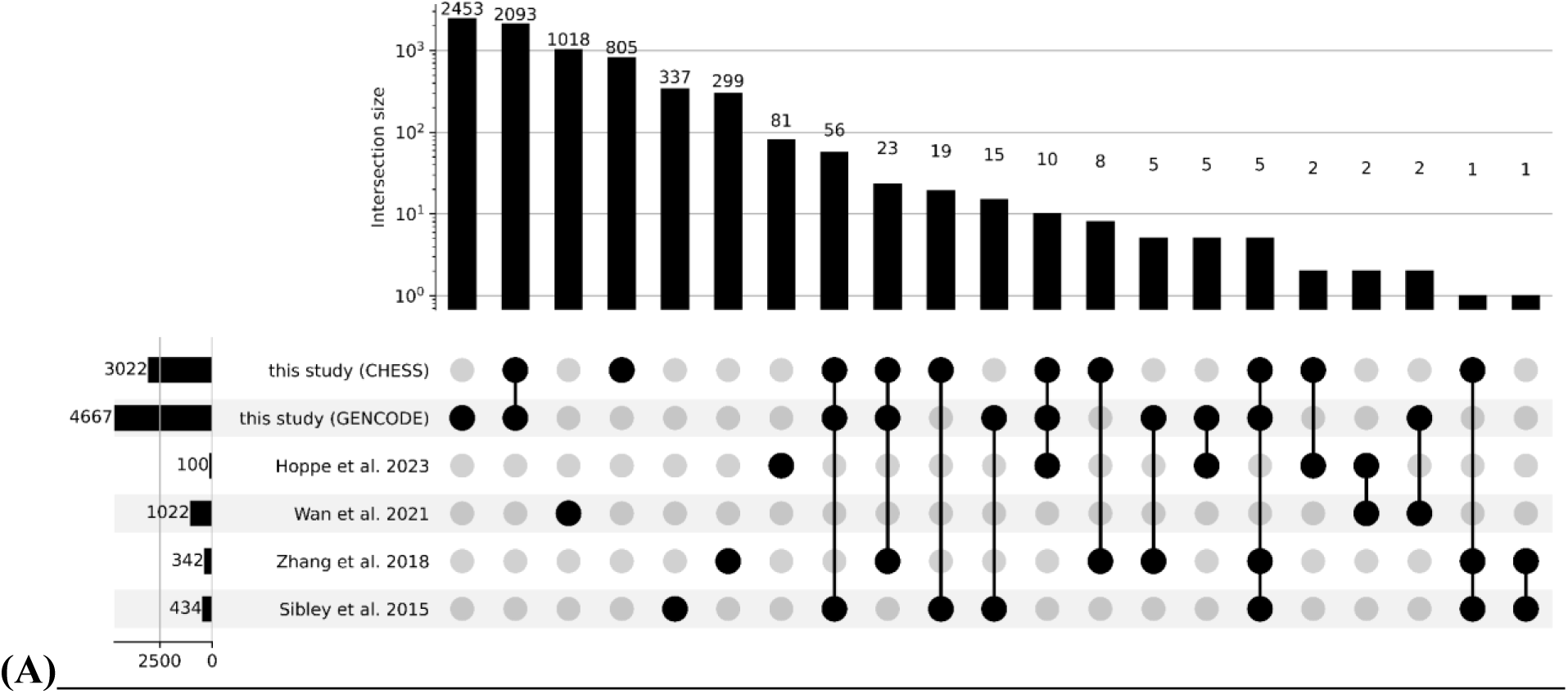
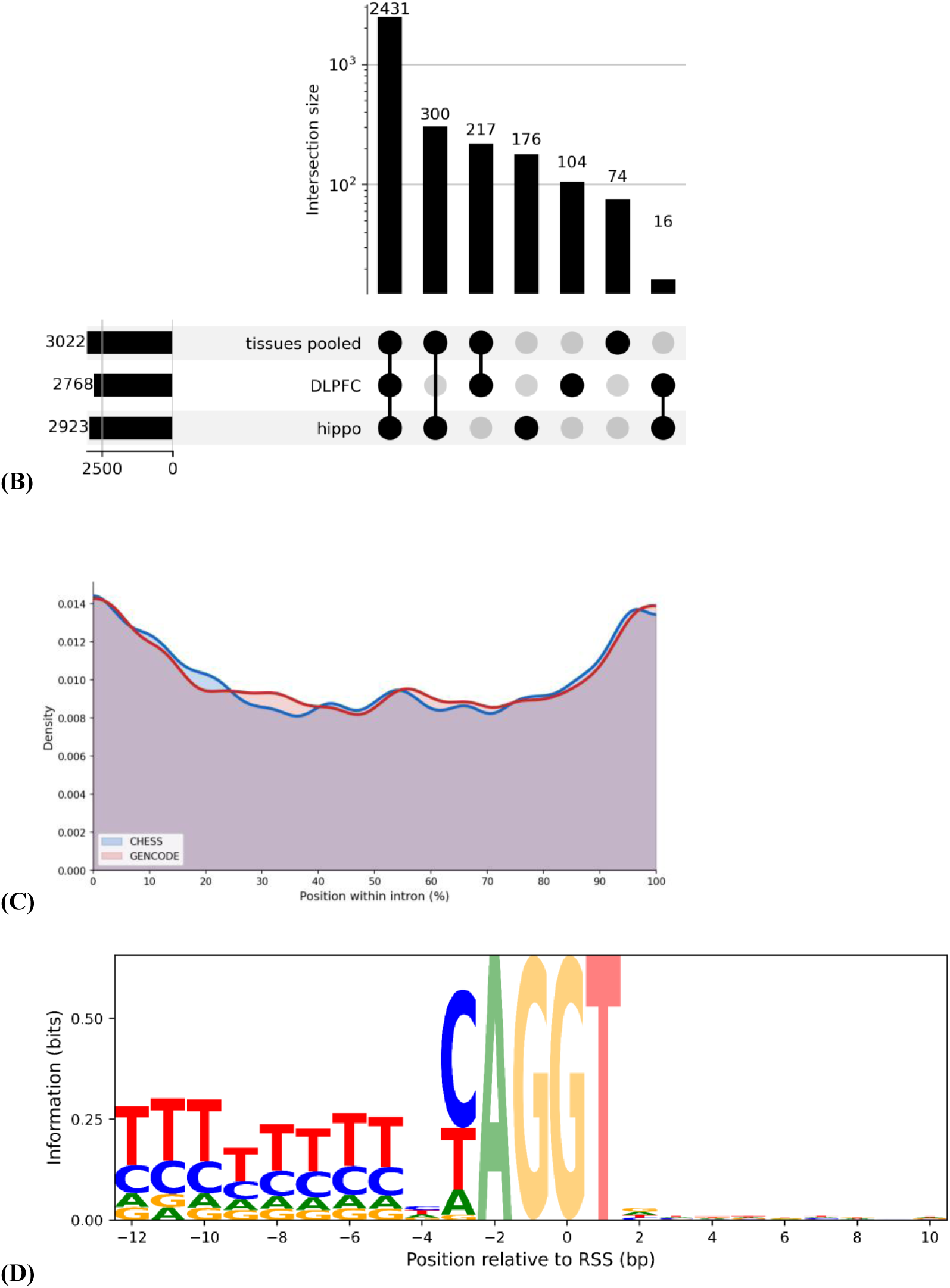
**(A)** UpSet plot (18) showing intersections of six sets of RSSs. This plot includes results from running our RSS detection pipeline on both CHESS and GENCODE, and it compares those sets two four prior RSS annotations (Hoppe, Wan, Zhang, Sibley). **(B)** UpSet plot comparing RSS sets that we obtain by pooling the two tissues versus analyzing samples from each one independently. **(C)** Kernel density estimation plot of relative locations of RSSs within the maximal introns that contain them. With either gene annotation, RSSs are slightly enriched near the edges of their introns. **(D)** Sequence logo shows the information content in sequence surrounding RSSs detected with CHESS and pooled tissues. By construction, our RSSs have 100% conservation at the core tetranucleotide `AGGT`. 86.7% of our RSSs feature a pyrimidine immediately upstream of the `AG`.

Only 124 (4.1%) of our 3,022 finalist RSSs overlap with any of the four previously published sets of RSSs. Using the more expansive GENCODE annotation, our method found many more RSSs (4,667), but still only 121 (2.6%) of those overlapped previously reported RSSs. One driver of RSS annotation disagreement is choice of underlying transcriptome annotation (see the CHESS vs. GENCODE rows of **Fig. 2A**). Furthermore, Hoppe *et al*. (6) only consider introns that appear constitutively; *i.e*., in all transcripts of a gene in their customized annotation.

Applying such a conservative restriction with GENCODE v47 would remove all but 4 of Hoppe *et al*.’s 100 RS introns. When we attempted to replicate Hoppe *et al*.’s (6) analysis using our Ribo-Zero data and GENCODE v47 filtered to allow introns that appear in ≥50% of the transcripts of a given gene, we found 83 RSS candidates, but only one was contained in the 100 from the original study, and none overlapped with our 3,022 finalists.

Using the CHESS annotation (v3.1.3) (15) as our reference, our 3,022 RSSs were found in 2,775 introns, some of which contained multiple RSSs (Table S1). 2,891 of our RSSs occur in protein-coding genes, 64 in long non-coding RNAs (lncRNAs), and 67 in transcribed pseudogenes. We replicated our analysis using the GENCODE annotation (v47) (16), which contains many more transcripts than either CHESS or RefSeq (Table S2). This replication yielded 4,667 RSSs, of which 2,093 overlapped those found when using CHESS (**Fig. 2A**) (Table S3).

**Table 1:** Total numbers of recursive splice sites (RSSs) found when using either the CHESS or GENCODE human annotation databases. Unlike the UpSet plots in **Fig. 2**, this table does not include prior RSS annotations; *e.g.*, the figure of 2,187 RSSs in this table is the sum of the sizes of all columns in **Fig. 2a** that include both RSS sets from this study (2,093+56+23+10+5=2,187).

| **Table 1:** Total numbers of recursive splice sites (RSSs) found when using either the CHES or GENCODE human annotation databases. Unlike the UpSet plots in **Fig. 2**, this table does not include prior RSS annotations; *e.g.*, the figure of 2,187 RSSs in this table is the sum of the sizes of all columns in **Fig. 2a** that include both RSS sets from this study (2,093+56+23+10+5=2,187). |  |  |  |
| --- | --- | --- | --- |
|  | **protein-coding genes** | **non-coding genes** | **all** |
| **CHES RSSs** | 2, 891 | 131 (4%) | 3, 022 |
| **GENCODE RSSs** | 4, 417 | 250 (5%) | 4, 667 |
| **CHES-only RSSs** | 752 | 83 (10%) | 835 |
| **GENCODE-only RSSs** | 2, 280 | 200 (8%) | 2, 480 |
| **RSSs in both** | 2, 137 | 50 (2%) | 2, 187 |
| **all** | 5, 169 | 333 (6%) | 5, 502 |

Although the numbers of RSSs found using the two annotation databases are different, 4,264/4,667 (91%) of our GENCODE RSSs are covered by introns in CHESS, and 2,977/3,022 (99%) of our CHESS RSSs are covered by introns in GENCODE. The explanation for most of the additional RSSs found when using GENCODE is that GENCODE contains many more alternative splice isoforms, which in turn generate many more RS pre-RNA targets. We also note that 2,601 (86.1%) of our CHESS-derived RS introns are shared by the MANE annotation (v1.5), which contains one carefully chosen transcript for each human protein-coding gene (19). The robustness of our method under changes to sample size is shown via resampling analysis in Table S6.

### Corroboration using spliced alignment to the genome

To corroborate the recursive splice sites found using 150bp target sequences (the semifinalist set), we used HISAT2 (20) to identify spliced alignments of the same reads to the full genome (see Methods). We observed that the number of reads aligning to our custom-designed targets was highly correlated with the number of reads that had spliced alignments to the genome (**Fig. 3A**). Ideally, the same reads from the Ribo-Zero data would align to both the targets (end-to-end) and the corresponding genomic locus (as a spliced alignment). As the figure shows, we found strong but not perfect correspondence between the two methods of identifying RSSs. In particular, 165 of the semifinalists for CHESS and 498 for GENCODE had no alignments supporting them from whole-genome spliced alignment, so we removed those to produce our final sets of RSSs.

**Figure 3:**
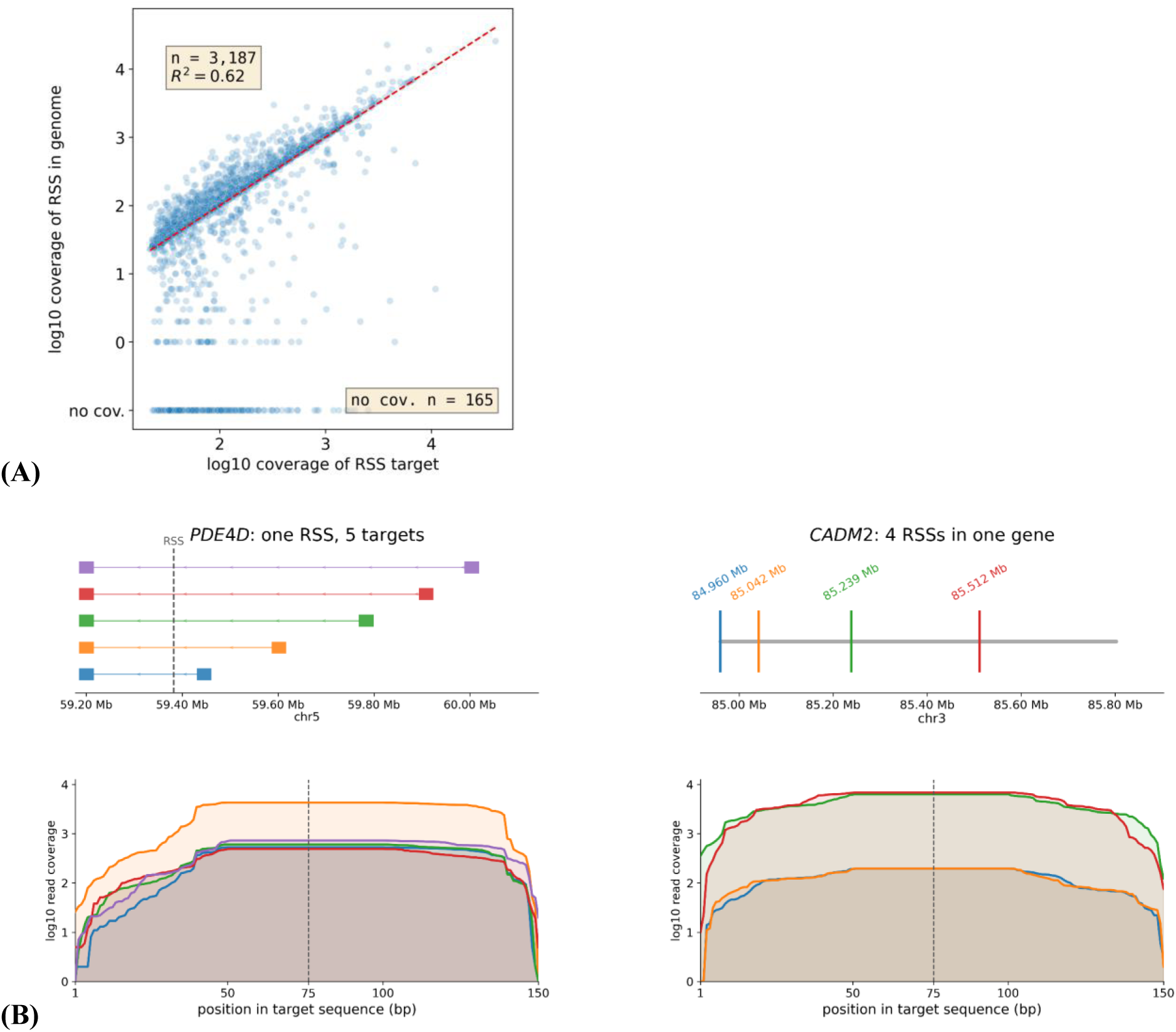
RNA-seq read coverage of finalist targets forms credible patterns. **(A)** Log–log scatterplot shows high concordance (*R*^2^ = 0.62) between unspliced alignment of Ribo-Zero reads to the candidate recursive splice sites’ (RSSs’) pre-RNA targets (horizontal axis) and spliced alignment to the genome (vertical axis). Each point shows a semifinalist RSS. Red line shows **`y = x`**. **(B)** Representative examples of read pileups to RS pre-RNA targets. On the left is *CADM2*, which contains four RSSs. On the right is a single RSS in *PDE4D* that, due to transcript diversity, appears in five targets with distinct upstream exons. Because the 150 bp targets were centered on putative RSSs and the reads were 100 bp long, all of the read coverage plots exhibit a plateau in the window ±25 bp around the RSS.

### Recursive splice sites tend to appear in longer introns

In **Fig. 4**, we show that the RS introns identified in this study have a median length of 10,114 bp, which is longer than the 2,339 bp and 2,110 median lengths found in introns from the GENCODE and CHESS annotations, respectively. To compute the median intron length in CHESS, we only considered introns longer than 200 bp, since our method only looked for RS sites in introns with ≥200 bp. To compute the distributions of RS intron lengths for each RSS annotation, we associated each RSS with the longest intron in CHESS that contains it and the surrounding ±50 bp window; if an RSS fit into another RSS’s intron, then we did not add another intron’s length to the distribution. (This latter counting method would yield significantly shorter intron lengths than those reported in prior papers that focus on long introns.) Under any annotation, RS intron lengths follow a unimodal, roughly log-normal, distribution. We found that RS introns are long, but not extremely long; our median length of ≈10 kbp is longer than Hoppe *et al.*’s (≈4 kbp) (6), but much shorter than Zhang *et al.*’s (≈36,837 kbp) (4) and Sibley *et al.*’s (≈17 kbp) (3). We report all CHESS-annotated introns, not just the longest, that contain our RSSs in Table S5.

**Fig. 4:**
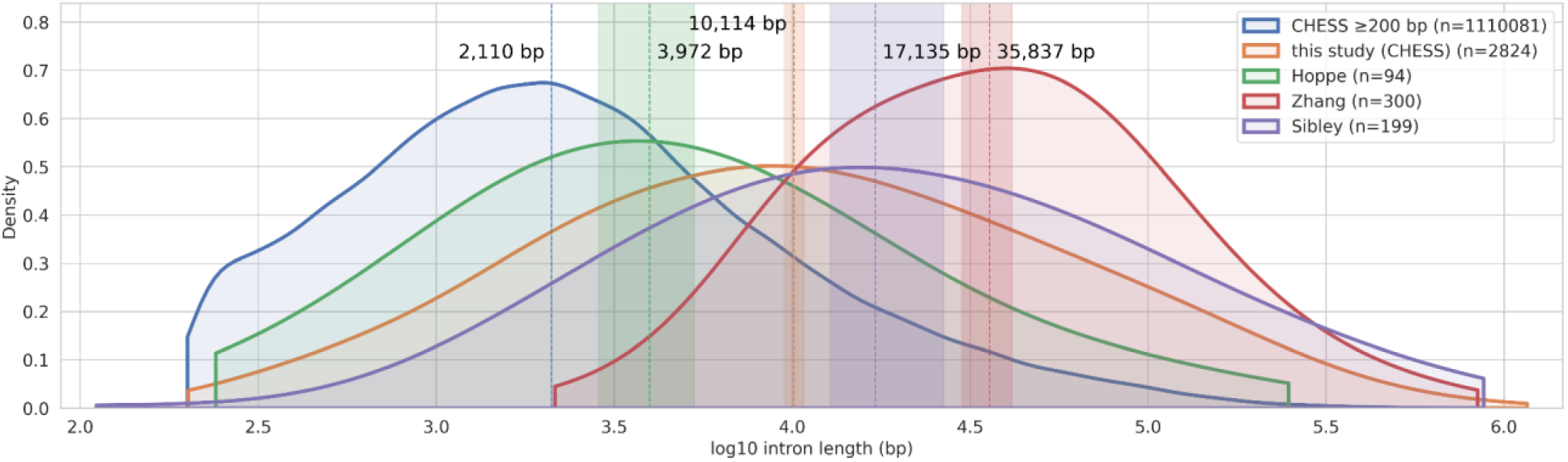
Kernel density estimation (KDE) plots of distributions of intron lengths found in several studies of recursive splicing. “CHESS ≥200 bp” includes all introns with length ≥200 bp in the CHESS annotation; “this study (CHESS)” refers to RS introns detected when running our method with CHESS; “Hoppe” includes the 94 RS introns covering 100 RSSs reported by Hoppe *et al*. 2023; “Zhang” refers to 300 RS introns covering 342 “candidate” RSSs reported by Zhang *et al*. 2018; and “Sibley” are the 199 RS introns covering 434 “putative” RS introns reported by Sibley *et al*. 2015. Covering introns were chosen as the longest introns in CHESS that contain a ±50 bp window around RSSs. Vertical dashed lines and text labels show medians, and surrounding bands show 95% bootstrap CI’s. Our GENCODE-based analysis (not shown) yielded a curve that was nearly identical to the CHESS-based one.

### RSSs are distinguished by RNA-binding protein motifs

We explored the question of whether recursive splicing is influenced by *cis* RNA-binding factors by testing whether specific RNA-binding protein (RBP) recognition sequences are enriched or depleted near RSSs. We queried the ATtRACT database of RBP binding motifs (21) in the intronic sequences ±100 bp of our RSSs, contrasted with a background of ±100 bp sequences around our 100,000 negative control loci. We found that sequences surrounding RSSs are enriched for binding motifs for PCBP1, PCBP2, and SRSF1, and are depleted for a binding motif for RBM8A (**Figure 5**). RBM8A is an end junction complex (EJC) component, so its significant depletion may indicate that the process of recursive splicing involves the downregulation of splicing inhibitors. (22) Statistics for all ATtRACT motifs are recorded in Table S4.

**Figure 5:**
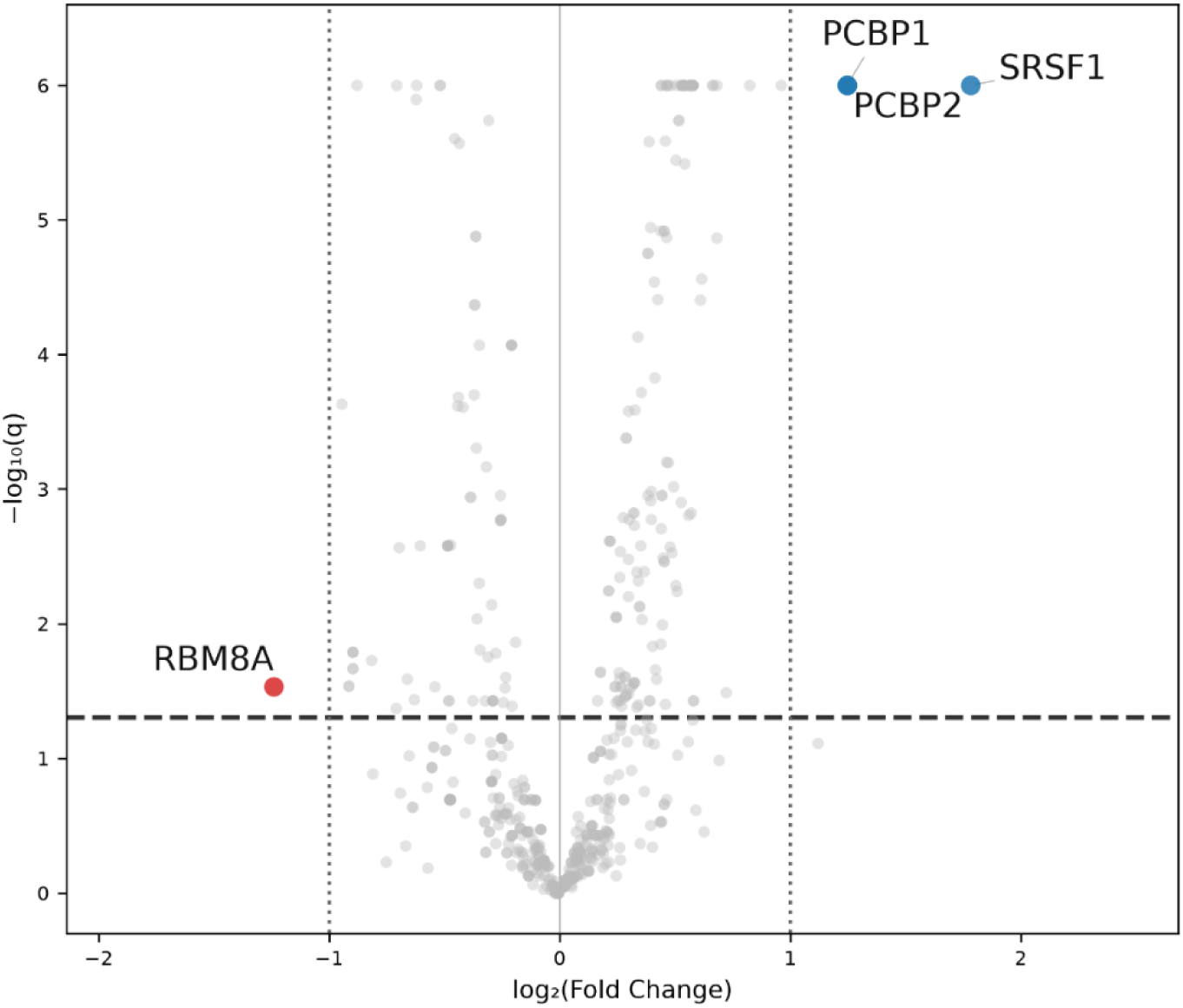
Volcano plots show enrichment and depletion of RBP-binding motifs in windows surrounding RSSs. Each point is a motif from the ATtRACT database (n=1,194). The horizontal axis shows base-2 logarithm fold change in motif abundance in the ±100 bp window around finalist RSSs versus negative controls. The vertical axis shows Benjamini–Hochberg-adjusted *p*-values from a two-sided Fisher’s exact test. For ease of visualization, low adjusted *p*-values were rounded up to 10^-6^. Under our thresholds of *p*_adj_ < 0.05 and |FC| > 2, a motif for RBM8A (red) is significantly depleted near RSSs, while motifs for PCBP1, PCBP2, and SRSF1 (blue) are significantly enriched.

## Discussion

We developed a novel method for detecting recursive splicing (RS) in human transcripts, finding many more RS sites than previously reported. Our alignments of RNA-seq reads to short, custom-designed pre-RNA targets were validated by spliced alignment of those same reads to the genome, indicating that reliable detection of RS events can be performed via focused analysis of small substrings from the transcriptome. Our results depended moderately on choice of annotation (CHESS v3 vs. GENCODE v47), although approximately two-thirds of the RS events found when using CHESS were also found with GENCODE.

Our detection of 3,022 RSSs represents a roughly 30-fold increase over the largest prior curated set (6). 4.1% of our RSSs overlap with prior reports, if we include “candidate” and “putative” RSSs from Zhang *et al.* (4) and Sibley *et al.* (3), respectively. Lariat enrichment (6), pulse-chase sequencing (5), 4sUDRB-Seq (4), and coverage-based methods (3) each capture slightly different features of the RS process, and we see our method as a more general approach that has the advantage of using RNA-seq data, which is a widely available data type.

Efforts to annotate RSSs are, arguably, at odds with Wan *et al.*’s (5) notion of stochastic splice site selection, in which the spliceosome component U2AF binds very frequently in nascent introns, creating RSSs that only slightly favor standard splicing motifs. Our method retains the classical view of splicing by only considering potential sites at canonical `AGGT` motifs, which occur within only 19% (1,022 of 5,469) of the RSSs reported by Wan *et al*.

Our results suggest that recursive splicing is a much more common process than was previously believed. If so, additional questions about RS need to be explored, including: does recursive splicing have a purpose beyond being an alternative mechanism to remove introns? And is recursive splicing a mechanism that evolved to assist in the removal of longer introns, or is it better understand as part of the standard splicing process? Further analysis, both from bulk sequence data and from targeted molecular methods, will be necessary to establish more reliable tools for, and a consensus annotation of, human recursive splicing events.

## Materials and Methods

### Genomic filtering for recursive splice site candidates

Our process for identifying recursively spliced (RS) introns began by searching through all annotated introns on all human chromosomes. We included every intron with length of at least 200 bp, under the assumption that the sterics of RS are unlikely to work with shorter introns. We required the internal splice site to consist of an acceptor site dinucleotide `AG` followed immediately by a donor site `GT`; i.e., every RS site must be centered on the 4-base sequence `AGGT`. We define a “subintron” as one of the two “halves” of an RS intron, separated by the RS site. Note that a single intron can have multiple RS sites. We then identified every potential RS site motif whose center lies at least 50 bp from either end of its intron.

We then used SPLAM, a deep learning-based tool for intron identification that has a low false positive rate (17), to score the two candidate subintrons defined by each RS site. SPLAM returns a score between 0 and 1 for each end of an intron (donor and acceptor), and we filtered our candidate set to retain only those RS introns that had a SPLAM score ≥0.5 for both the acceptor of the upstream subintron and the donor of the downstream subintron; these are the internal splice termini that comprise the `AGGT` motif. Filtering with SPLAM reduced our candidate set by 90%, from 22,378,870 subintron pairs to 2,219,760. Note that we considered intron–RS site pairs (or equivalently, pairs of subintrons) at this stage of our analysis in order to track both the RS site and the intron in which it appears; some of the introns of interest were overlapping in the genome.

One source of false positives is loci that are “falsely” intronic because they are entirely within an intron of one transcript, but are at an intron–exon junction of another transcript. To avoid this source of error, we removed candidate loci that fell at intron–exon junctions in any transcript, with two exceptions: “mutually exclusive” loci for which we can construct a target with an upstream exon that never co-occurs in a transcript with the RSS’s downstream exon, and “short” loci, for which the RSS’s downstream exon is shorter than our minimum alignment anchor length of 25bp. These two categories accounted for 212 of our 3,022 finalist RSSs.

### Searching RNA-seq reads

To identify experimental evidence of recursive splicing, we needed RNA sequences that included partially spliced introns. For this reason, we used total RNA-seq experimental data, which sequences all of the mRNA in a tissue sample (including pre-mRNA) rather than just polyadenylated transcripts, collected using a protocol (“Ribo-Zero Gold”) that removes ribosomal RNA. Our primary data source was 900 previously published samples collected by the Lieber Institute for Brain Development’s (LIBD) BrainSeq Phase II project (14) from postmortem dorsolateral prefrontal cortex (DLPFC) and hippocampus samples. In total, the 900 samples included 101,928,170,464 paired-end reads, where all reads had a length of 100 bp. To avoid alignment issues with short transcript fragments, we treated paired reads as if they were unpaired in our analysis.

We first filtered the reads by removing those that aligned to the mature transcriptome (i.e., mRNA transcripts from which introns were removed) extracted from the CHESS v3.1.3 human annotation (15), excluding pseudogenes. We aligned reads using Bowtie2 (23) with the parameters `--very-fast -k 1`. This step left 47,298,075,084 reads that did not align. Separately, we aligned the full set of reads to the processed transcriptome from GENCODE v47 (Mudge et al. 2025), excluding transcripts whose biotypes contain “decay”, “pseudogene”, “intron”, or “TEC,” which left 54,630,095,380 reads. We then filtered again by removing reads that Bowtie2 (`--very-fast -k 1`) successfully aligned to the genome (GRCh38.p14). This step left 8,723,414,672 reads following CHESS-based filtering and 8,737,777,756 reads following GENCODE-based filtering (∼9% of the initial read set). Because Bowtie2 is an unspliced aligner, these two filtering steps controlled for false positives by removing reads that did not derive from the premature transcripts that we were seeking. In this context, “premature” is synonymous with “not fully spliced” or “intermediate isoform reads” (as in (24)).

### Construction of RS pre-RNA targets

We then constructed pre-mRNA alignment targets consisting of 150 bp subsequences unique to each candidate recursive splice site (RSS) in each annotated transcript from CHESS v3.1.3 or GENCODE v47. Each target represented a partially spliced transcript that had all introns excised up to and including its upstream subintron, but still retained all downstream introns, including the downstream subintron. Each target consisted of 75 bp of upstream flanking exonic sequence concatenated with 75 bp downstream of the RSS in a transcript (**Fig. 1B**). Since most transcripts of the same gene will yield identical RS pre-mRNA targets, and some human genes have near-identical paralogs, we de-duplicated target sequences, retaining only one distinct copy of each. For each target, we kept track of (i) the annotated intron from which it derived, (ii) the candidate RSS, and (iii) the set of transcripts (one or more) that it might appear in. In ∼10% of cases (241,902 / 2,508,642), the flanking sequence was too short (i.e., the RS intron followed <75 bp of upstream exonic sequence, or the RS site was <75 bp away from the 3′ end of the gene). In those cases, we simply constructed a shorter RS-preRNA target from the maximal useful sequence that was available. Our RS pre-mRNA targets had a mean length of 149.37 for CHESS and 147.71 for GENCODE.

### Negative controls

We constructed a negative control set by uniformly sampling 100,000 non-overlapping introns from CHESS v3.1.3 with lengths ≥200 bp. For each intron, we uniformly sampled an RS site that was at least 50 bp from either end, while ensuring that the center of the control sequence did not match the RS motifs `AGGT` or `AGGC` (`GC` is the second-most common human 5′ donor dinucleotide). We used this negative control set to empirically set thresholds at each stage of filtering in our pipeline, prioritizing a low false discovery rate. Although there is a small chance that non-canonical recursive splicing might occur at any of these loci, the vast majority are expected to be negative. We used these negative controls to set thresholds yielding a false discovery rate of 0.01%.

### Data acquisition from prior recursive splice site annotations

We obtained 100 RSSs from Hoppe *et al.* (6) and 342 candidate RSSs from Zhang *et al.* (4). We collected 434 “putative” distinct RSSs from Table S1 of Sibley et al., but we note that their most conservative RSS annotation contains just 9 introns. We also obtained 5,469 RSSs from Table S4 of Wan *et al.* (5), which we filtered to 1,022 by keeping only those with `AGGT` motifs. All of these sets used the GRCh37 coordinate system, which we converted to GRCh38 using a public Python package (https://github.com/jeremymcrae/liftover/).

### Searching for significant enrichment/depletion of motifs

We query the ATtRACT database of RNA-binding protein motifs with the MEME suite tool FIMO (`--text --thresh 1e-4`) (25). *p*-values for motif hits were computed with a two-sided Fisher’s exact test and adjusted with the Benjamini–Hochberg procedure.

### *De novo* gene annotation with StringTie3

For *de novo* gene annotation, we performed paired spliced alignment of all 900 Ribo-Zero samples to GRCh38 with HISAT2 (20) using `--no-softclip --fast --max- intronlen 1240120`. (The longest intron in GENCODE is 1,240,120 bp.) We input these 900 alignment BAMs into StringTie3 with the `-N` parameter, which optimizes the system for assembly from total RNA-seq samples (9).

### Code and data availability

The code for the project is stored in a publicly available GitHub repository, https://github.com/DavidB256/recursive_introns. Code instructions are found in a Snakemake (26) pipeline (see file [snakefile](https://github.com/DavidB256/recursive_introns/blob/main/snakefile)) or in a Jupyter notebook (27) in the repository. FTP addresses for the data downloaded from other sources can be found at the top of the [snakefile](https://github.com/DavidB256/recursive_introns/blob/main/snakefile). For statistical analyses, we used the Python libraries SciPy (28) and statsmodels (29). Portions of our code base were written and organized by Anthropic LLMs via Claude Code.

## Acknowledgements

The authors thank Rajiv McCoy and Erik Andersen for helpful comments on earlier drafts of this paper. This work was supported in part by the U.S. National Institutes of Health under grants R01-HG006677 and R35-GM130151.

## Supporting information captions

All coordinates in our supplemental tables are 1-based in GRCh38.\

**Table S1:** Our semifinalist and finalist recursive splice sites, as determined from CHESS v3.1.3. Columns: chrom, locus, strand, gene (gene name), gene_type, exonic_type (whether the RSS is at an intron–exon junction, and the type of junction if it is at one), is_finalist (whether it is a “finalist”, or just a “semifinalist”), containing_transcripts (semicolon-separated list of CHESS transcript IDs of transcripts that include the RSS), target_coverage (number of reads that align to the 150 bp target), spliced_coverage (number of reads that align spliced over an upstream subintron created by the RSS), intron_start (start locus of longest intron that contains RSS), intron_end (end locus of longest intron that contains RSS), intron_length (length of longest intron that contains RSS)

**Table S2:** Our semifinalist and finalist recursive splice sites, as determined from GENCODE v47. Contains same columns as Table S1, through spliced_coverage.

**Table S3:** Overlap of RSSs among different annotations. Columns: chrom, strand, locus, in_this_study_CHESS, in_this_study_GENCODE, in_Hoppe_2023, in_Zhang_2018, in_Sibley_2015, in_Wan_2021

**Table S4:** FIMO motif enrichment results from ATtRACT RNA-binding protein (RBP) motif database near finalist CHESS RSSs. Columns: motif_id, rbp_gene, consensus_seq (of motif), log2fc (fold-change between RSSs and negative controls), p (*p*-value from two-sided Fisher’s exact test), q (BH-adjusted *p*-value), enrichment (NS=not significant), hits_rss (count of motif near RSS), n_rss (count of RSSs used), hits_neg (count of motif near negative control loci), n_neg (count of negative control loci used)

**Table S5:** Introns in CHESS that cover at least one finalist RSSs from CHESS. Columns: chrom, intron_start, intron_end, strand, intron_length, rss_coordinates (semicolon-separated list of coordinates of RSSs that appear within the intron, at least 50 bp from either end)

**Table S6:** RSSs captured from CHESS annotation on subsampled RNA-seq data. Columns: read_fraction_pct (proportion of Ribo-Zero RNA-seq reads kept in subsample), n_semifinalists (number of semifinalist RSSs captured), n_finalists (number of finalist RSSs captured)

## References

1. Hatton AR, Subramaniam V, Lopez AJ. Generation of Alternative *Ultrabithorax* Isoforms and Stepwise Removal of a Large Intron by Resplicing at Exon–Exon Junctions. Mol Cell. 1998 Dec 1;2(6):787–96. doi:10.1016/S1097-2765(00)80293-2

2. Duff MO, Olson S, Wei X, Garrett SC, Osman A, Bolisetty M, et al. Genome-wide identification of zero nucleotide recursive splicing in Drosophila. Nature. 2015 May;521(7552):376–9. doi:10.1038/nature14475

3. Sibley CR, Emmett W, Blazquez L, Faro A, Haberman N, Briese M, et al. Recursive splicing in long vertebrate genes. Nature. 2015 May;521(7552):371–5. doi:10.1038/nature14466

4. Zhang XO, Fu Y, Mou H, Xue W, Weng Z. The temporal landscape of recursive splicing during Pol II transcription elongation in human cells. PLOS Genet. 2018 Aug 27;14(8):e1007579. doi:10.1371/journal.pgen.1007579

5. Wan Y, Anastasakis DG, Rodriguez J, Palangat M, Gudla P, Zaki G, et al. Dynamic imaging of nascent RNA reveals general principles of transcription dynamics and stochastic splice site selection. Cell. 2021 May 27;184(11):2878–2895.e20. doi:10.1016/j.cell.2021.04.012 PubMed PMID: 33979654.

6. Hoppe ER, Udy DB, Bradley RK. Recursive splicing discovery using lariats in total RNA sequencing. Life Sci Alliance. 2023 Jul 1;6(7). doi:10.26508/lsa.202201889 PubMed PMID: 37137707.

7. Joseph B, Kondo S, Lai EC. Short cryptic exons mediate recursive splicing in Drosophila. Nat Struct Mol Biol. 2018 May;25(5):365–71. doi:10.1038/s41594-018-0052-6

8. Mueller RL, Adams AN. Recursive splicing—a mechanism of intron removal with an unexplored role in the largest genomes. J Mol Evol. 2025 Aug 1;93(4):474–7. doi:10.1007/s00239-025-10261-9

9. Shinder I, Pertea G, Hu R, Rudnick Z, Pertea M. StringTie3 improves total RNA-seq assembly by resolving nascent and mature transcripts. Nat Methods. 2026 Jun;23(6):1126–37. doi:10.1038/s41592-026-03080-3

10. Taggart AJ, DeSimone AM, Shih JS, Filloux ME, Fairbrother WG. Large-scale mapping of branchpoints in human pre-mRNA transcripts in vivo. Nat Struct Mol Biol. 2012 Jul;19(7):719–21. doi:10.1038/nsmb.2327

11. Pineda JMB, Bradley RK. Most human introns are recognized via multiple and tissue- specific branchpoints. Genes Dev. 2018 Apr 1;32(7–8):577–91. doi:10.1101/gad.312058.118 PubMed PMID: 29666160.

12. Adusumalli S, Ngian ZK, Lin WQ, Benoukraf T, Ong CT. Increased intron retention is a post-transcriptional signature associated with progressive aging and Alzheimer’s disease. Aging Cell. 2019;18(3):e12928. doi:10.1111/acel.12928

13. Ling JP, Pletnikova O, Troncoso JC, Wong PC. TDP-43 repression of nonconserved cryptic exons is compromised in ALS-FTD. Science. 2015 Aug 7;349(6248):650–5. doi:10.1126/science.aab0983

14. Collado-Torres L, Burke EE, Peterson A, Shin J, Straub RE, Rajpurohit A, et al. Regional Heterogeneity in Gene Expression, Regulation, and Coherence in the Frontal Cortex and Hippocampus across Development and Schizophrenia. Neuron. 2019 Jul 17;103(2):203–216.e8. doi:10.1016/j.neuron.2019.05.013 PubMed PMID: 31174959.

15. Varabyou A, Sommer MJ, Erdogdu B, Shinder I, Minkin I, Chao KH, et al. CHESS 3: an improved, comprehensive catalog of human genes and transcripts based on large-scale expression data, phylogenetic analysis, and protein structure. Genome Biol. 2023 Oct 30;24(1):249. doi:10.1186/s13059-023-03088-4

16. Mudge JM, Carbonell-Sala S, Diekhans M, Martinez JG, Hunt T, Jungreis I, et al. GENCODE 2025: reference gene annotation for human and mouse. Nucleic Acids Res. 2025 Jan 6;53(D1):D966–75. doi:10.1093/nar/gkae1078

17. Chao KH, Mao A, Salzberg SL, Pertea M. Splam: a deep-learning-based splice site predictor that improves spliced alignments. Genome Biol. 2024 Sep 16;25(1):243. doi:10.1186/s13059-024-03379-4

18. Lex A, Gehlenborg N, Strobelt H, Vuillemot R, Pfister H. UpSet: Visualization of Intersecting Sets. IEEE Trans Vis Comput Graph. 2014 Dec;20(12):1983–92. doi:10.1109/TVCG.2014.2346248

19. Morales J, Pujar S, Loveland JE, Astashyn A, Bennett R, Berry A, et al. A joint NCBI and EMBL-EBI transcript set for clinical genomics and research. Nature. 2022 Apr;604(7905):310–5. doi:10.1038/s41586-022-04558-8

20. Kim D, Paggi JM, Park C, Bennett C, Salzberg SL. Graph-based genome alignment and genotyping with HISAT2 and HISAT-genotype. Nat Biotechnol. 2019 Aug;37(8):907–15. doi:10.1038/s41587-019-0201-4

21. Giudice G, Sánchez-Cabo F, Torroja C, Lara-Pezzi E. ATtRACT—a database of RNA-binding proteins and associated motifs. Database. 2016 Jan 1;2016:baw035. doi:10.1093/database/baw035

22. The UniProt Consortium. UniProt: the Universal Protein Knowledgebase in 2025. Nucleic Acids Res. 2025 Jan 6;53(D1):D609–17. doi:10.1093/nar/gkae1010

23. Langmead B, Salzberg SL. Fast gapped-read alignment with Bowtie 2. Nat Methods. 2012 Apr;9(4):357–9. doi:10.1038/nmeth.1923

24. Choquet K, Baxter-Koenigs AR, Dülk SL, Smalec BM, Rouskin S, Churchman LS. Pre-mRNA splicing order is predetermined and maintains splicing fidelity across multi-intronic transcripts. Nat Struct Mol Biol. 2023 Aug;30(8):1064–76. doi:10.1038/s41594-023-01035-2

25. Grant CE, Bailey TL, Noble WS. FIMO: scanning for occurrences of a given motif. Bioinformatics. 2011 Apr 1;27(7):1017–8. doi:10.1093/bioinformatics/btr064

26. Köster J, Rahmann S. Snakemake—a scalable bioinformatics workflow engine. Bioinformatics. 2012 Oct 1;28(19):2520–2. doi:10.1093/bioinformatics/bts480

27. Granger BE, Pérez F. Jupyter: Thinking and Storytelling With Code and Data. Comput Sci Eng. 2021 Mar;23(2):7–14. doi:10.1109/MCSE.2021.3059263

28. Virtanen P, Gommers R, Oliphant TE, Haberland M, Reddy T, Cournapeau D, et al. SciPy 1.0: fundamental algorithms for scientific computing in Python. Nat Methods. 2020 Mar;17(3):261–72. doi:10.1038/s41592-019-0686-2

29. Seabold S, Perktold J. Statsmodels: Econometric and Statistical Modeling with Python. In. Austin, Texas; 2010 [cited 2026 Mar 1]. p. 92–6. Available from: https://doi.curvenote.com/10.25080/Majora-92bf1922-011 doi:10.25080/Majora-92bf1922-011

